# Characterization of transcriptomic profiles underlying gross morphological changes observed in semelparous pink salmon (*Oncorhynchus gorbuscha*)

**DOI:** 10.64898/2026.02.12.705573

**Authors:** Max Butensky, Michael Phelps

## Abstract

Pacific salmon (*Oncorhynchus spp*.) undergo intricate physiological changes during maturation as they migrate to spawning beds and breed before succumbing to a programmed senescence (semelparous life cycle). Research into the physiological mechanisms of semelparity in salmon has identified a clear and progressive rise in sex and stress hormone levels throughout their migration, which correlates with the emergence of morphological traits, as well as changes in behavioral patterns. We examined transcriptional changes in three critical tissues (gonads, head kidney, and skeletal muscle) across the spawning migration in male and female Pink salmon (*Oncorhynchus gorbuscha*) to capture the molecular changes occurring in these tissues during maturation and senescence. Major transcriptional changes occurred around the time of spawning, while only modest transcriptional changes were found as the fish migrated between saltwater and freshwater. Examination of enriched biological pathways identified the signatures of semelparous catabolic processes in all tissues and a strong immune response in somatic tissues. Evidence of shifts in lipid energy mobilization were also seen in somatic tissues. A closer investigation of the expression patterns of endocrine hormone receptors showed that many endocrine pathways prioritized expression of specific dominant ohnologs to orchestrate much of the hormone response in the analyzed tissues. Our characterization of the transcriptional profiles in migrating pink salmon adds critical context to link the molecular changes occurring in tissues to the physiological transitions that define semelparous maturation in Pacific salmon.

**NEW & NOTEWORTHY:** Large transcriptional changes occurred in the gonads, head kidney, and skeletal muscle of pink salmon during the final stages of their spawning migration. Across the tissues and sexes, spawning was marked by coordinated activation of catabolic programs (autophagy, proteolysis, cell death), and a strong immune response in somatic tissues, alongside lipid mobilization. Endocrine receptor expression analyses revealed that the response to hormones was primarily mediated by a limited number of dominant ohnologs.

## 1. INTRODUCTION

The anadromous journey of Pacific salmon to return to natal spawning grounds involves coordinated physiological responses that result in dramatic physical transformations, sexual maturity, and death (semelparity). Semelparous salmon have one shot at breeding, and the maturation process is critical to ensuring that they reach the spawning ground and produce quality gametes before succumbing to their deteriorating condition. The hypothalamic-pituitary-gonadal (HPG) endocrine axis plays a crucial role in reproductive development through its regulation of gonadal maturation and sex steroid production, which has been associated with end-of-life phenotypes in Pacific salmon (1). A functional link between reproductive maturation and semelparity in salmon has previously been established in gonadectomized sockeye salmon, which did not exhibit semelparous death, or rapid emergence of sexual dimorphic traits (1). It was also found that the regeneration of excised gonads in these fish in future years correlated with senescence and the development of reproductive characteristics. These studies support the idea that gonadal maturation in semelparous salmon is a driver of progressive tissue degeneration and death, but how this process functions at a molecular level is unknown.

Sex steroids synthesized in the gonads contribute to phenotypic changes that emerge during reproductive maturation, and these endocrine changes can influence the fitness of salmon as they reach the spawning grounds (2–6). Reproductive success in salmon is locally adapted, with fish from different waterways exhibiting variation in their physiological changes and timing of spawning (7–10). Notably, during a salmon’s migration, there is a steady decline in skeletal muscle quality as they approach the spawning grounds, correlating with the use of that tissue as an energy source in the absence of feeding (11, 12). Activated catabolic processes in skeletal muscle break down energy stores to sustain the organism during migration and to ensure successful reproductive development of the gonads in time for spawning (13–15). The androgens have been shown to act as immunosuppressors in both males and females, impairing optimal microbial defense responses and potentially contributing to the prevalence of prespawn mortality as fish succumb to elevated microbial loads (4, 16, 17). Sex steroid production is hypothesized to induce hyperplasia of adrenal tissues, driving the observed rise in the synthesis of cortisol levels during the salmon migration (1, 18, 19). It is this overactive cortisol production that is hypothesized to be a major contributor to semelparous senescence (20, 21). Hypercorticotropism and the physiological changes associated with it are characteristic of semelparity across divergent taxa (e.g., insects, mollusks, fishes, mammals), indicating a clear phenotypic trend resulting in programmed death (12, 20, 22, 23).

To gain insight into the molecular drivers of reproductive maturation and semelparity in Pacific salmon, we characterized the transcriptomic landscape of the gonads, head kidney, and skeletal muscle tissue during the spawning migration in pink salmon (*Oncorhynchus gorbuscha*). The research examined novel, differentially regulated gene pathways associated with key phenotypic changes that occur during salmon maturation. This included a detailed analysis of gene expression patterns associated with known endocrine processes. By examining tissues that function as endocrine sources (e.g., gonads and head kidney) and sinks (skeletal muscle), we were able to gain insight into how stress and sex hormones might influence maturation in Pacific salmon. The findings of this study provide insight into the molecular mechanisms underlying maturation and semelparity in Pacific salmon, a taxonomic group with strong ecological, economic, and cultural significance.

## 2. Methods and Materials

### 2.1 Collection of Wild Pink Salmon Tissues and RNA Isolation

Tissue samples, in accordance with IACUC (#6607), were collected from wild odd-year pink salmon on their return migration to Washington State, USA. This was done under the scientific collection permit (WDFW: phelps21-218), scientific research permit (NOAA #25795), and recovery permit (USFW: PER0016959-0). We collected 3-5 individuals of each sex from three locations: (1) Washington State marine zone 8-2 (47.924162, -122.338837) on August 9th,2021, (2) the Skykomish River (9.5 river miles, 47.848968, -121.831456) on August 28th, 2021, and (3) spawning grounds on the Sultan River (Osprey Park, 47.870524, -121.825933) on September 26th 2021 (Figure 1). The Skykomish River sampling location was approximately 30 river miles from the Puget Sound and only approximately 2 river miles from the spawning ground sampling location on the Sultan River. Fish were caught with rod and reel, euthanized by blunt force followed by exsanguination, after which samples were immediately taken from the epaxial white skeletal muscle, head kidney, and testes. The samples were stored in RNAlater (ThermoFisher Scientific, cat. AM7021) at 4C for 24 hours, after which RNAlater was removed, and the samples were stored at -80C until processing. Total RNA was extracted from the tissues using TRIzol reagent (ThermoFisher, cat. 15596026), following the manufacturer’s protocols, and with the aid of 5PRIME Phase Lock heavy gel tubes (Quantabio, cat. 2302830), and 1-bromo-3-chloropropane (ThermoFisher, cat. 106860025) to aid in the isolation of RNA from DNA and protein. The total RNA was quantified on a Qubit fluorometer (ThermoFisher, cat. Q33238), and quality was checked using an Agilent 5200 fragment analyzer (Agilent, cat. M5310AA) at the WSU Laboratory for Biotechnology and Bioanalysis. Only samples with RNA Integrity Number greater than 6 were sent for sequencing.

**Figure 1.**
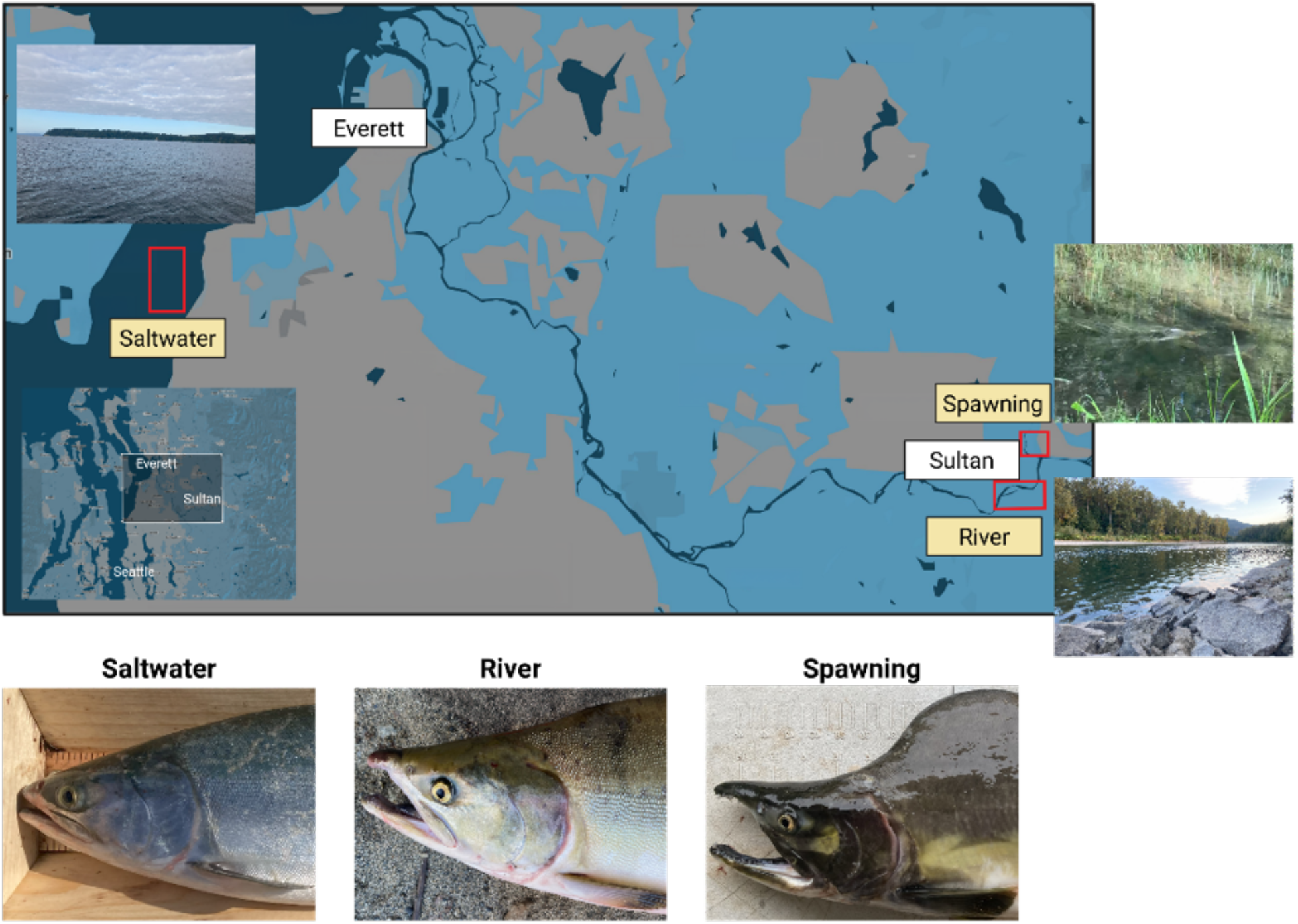
Sampling locations and physical transformation of pink salmon during their return migration. Pink salmon were collected from the Puget Sound (Saltwater), the Skykomish River (River), and spawning grounds on the Sultan River (Spawning) in Washington State, USA. Pink salmon undergo a significant physical transformation upon their entry into freshwater with a loss of silver coloration and the development of secondary sex traits upon spawning. Photos taken 3.5 weeks apart.

### 2.2 SEQUENCING and Expression Analysis

The total RNA was polyA enriched to capture mRNA transcripts and sequenced using an Illumina NovaSeq with paired-end 150 bp reads (Novogene). The raw reads (FASTQ) were filtered by sample, adapters were trimmed, and low-quality reads were removed using the default setting of TrimGalore package (version 0.6.4) ^6,24–26^. Next, cleaned reads were annotated and quantified against the transcriptome generated from *O. gorbuscha* reference transcriptome (OgorEven_v1.0) using the Salmon 1.8.0 package. Salmon leverages quasi-mapping to rapidly align reads to the transcriptome and then uses bias modeling to account for technical biases to accurately quantify the reads. This was first done by creating a decoy-aware index of the transcriptome and genome using a k-mer size of 31(24, 25). Using the salmon quant function and the validateMapping parameter, pair-end reads were quantified against the decoy-aware index to generate quantifications of captured mRNA for each sample. The Rstudio package tximport calculated transcript abundance, then normalized using vst function on the transcript-level estimates to correct for transcript length differences before compiled into a count matrix of 40,756 genes.

Differential expression analysis utilized DESeq2 package to identify relative changes in gene transcripts across each timepoint and biological replicate and further normalize across st (Figure 2)(25). Changes in the global gene transcription were quantified across the sequential stages of the pink salmon return migration by analyzing differential expression from saltwater to river and river to spawning ground. To gain insight into the pattern of gene expression changes, we categorized genes based on their changes between the migratory timepoints: *significantly* UP-regulated, DOWN-regulated, or genes with no significant change in expression (Figure 2). This criterion was collated to generate nine groups of genes that share in relative expression (Figure 2, Supplemental Table 1. Gene Ontology enrichment for each group identified biological pathways changing during migration by generating an index of *O. gorbuscha* peptide sequences homologs and associated gene ontology terms using the online portal of EggNOG mapper (26, 27). Next, enrichments for target genes were performed with the Rscript gene ontology Mann-Whitney U-test (GO_MWU) using fisher’s exact test for significance against the background of non-target genes (28). This tool seeks groupings of gene ontology terms across the global list of genes to identify anomalies from an even distribution and is thus valuable for novel models because it allows for the use of custom data as background (Figure 3). Terms were considered significant at a false discovery rate of 0.05, and a subset was presented.

**Figure 2:**
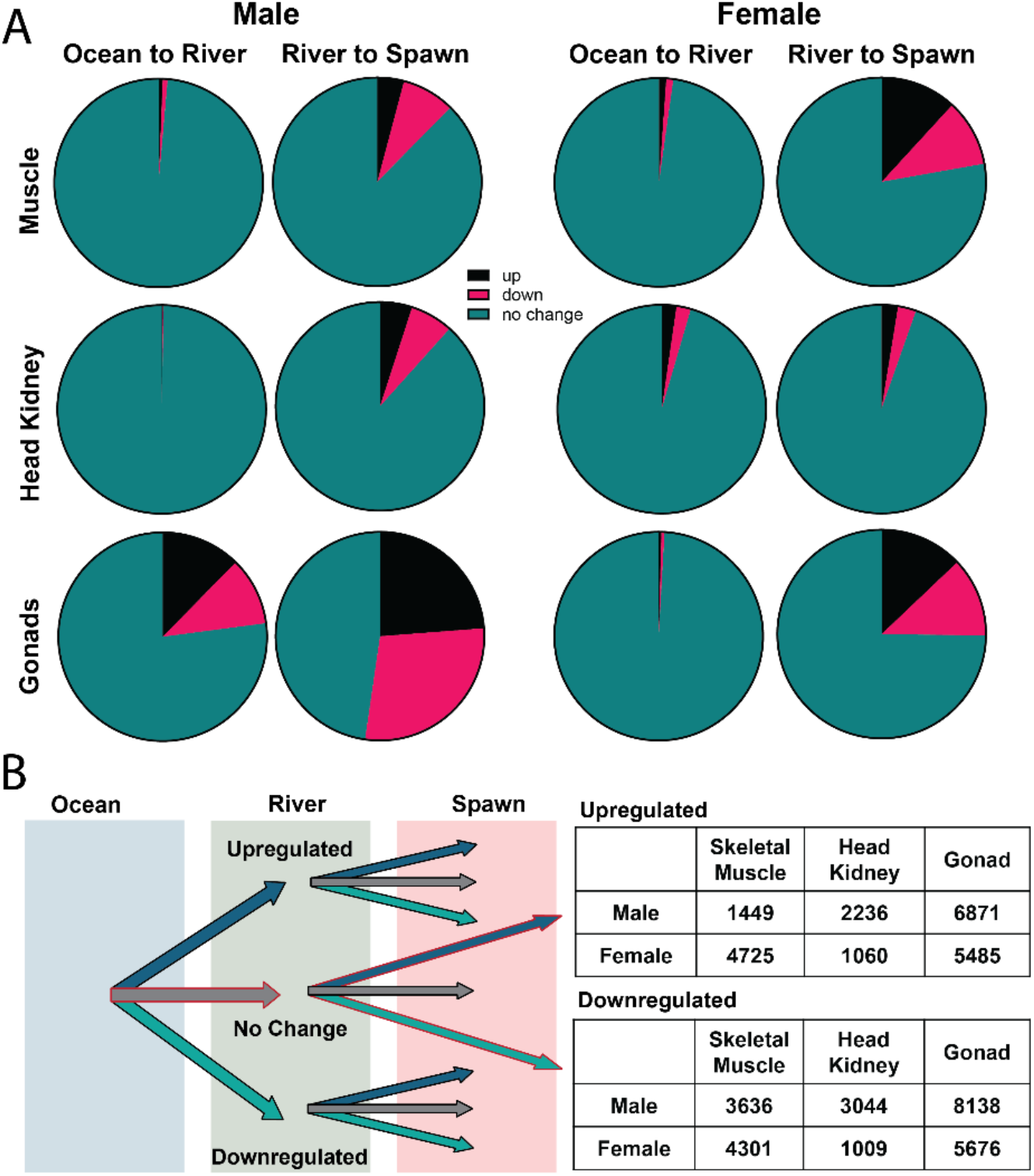
Patterns of differentially expression genes during the pink salmon migration. **(A)** Relative gene expression between tissues and sexes (up, down, no change), across 2 migratory transitions (Ocean to River and River to Spawning). **(B)** Filtration of genes by relative expression (+/-0 Log2foldchange, p-value < 0.05) across all stages generated 9 expression patterns, 2 of which were used for further analysis. Data shown for the number of genes that did not change significantly from ocean to river but were significantly up- or down-regulated from River to Spawn.

**Figure 3:**
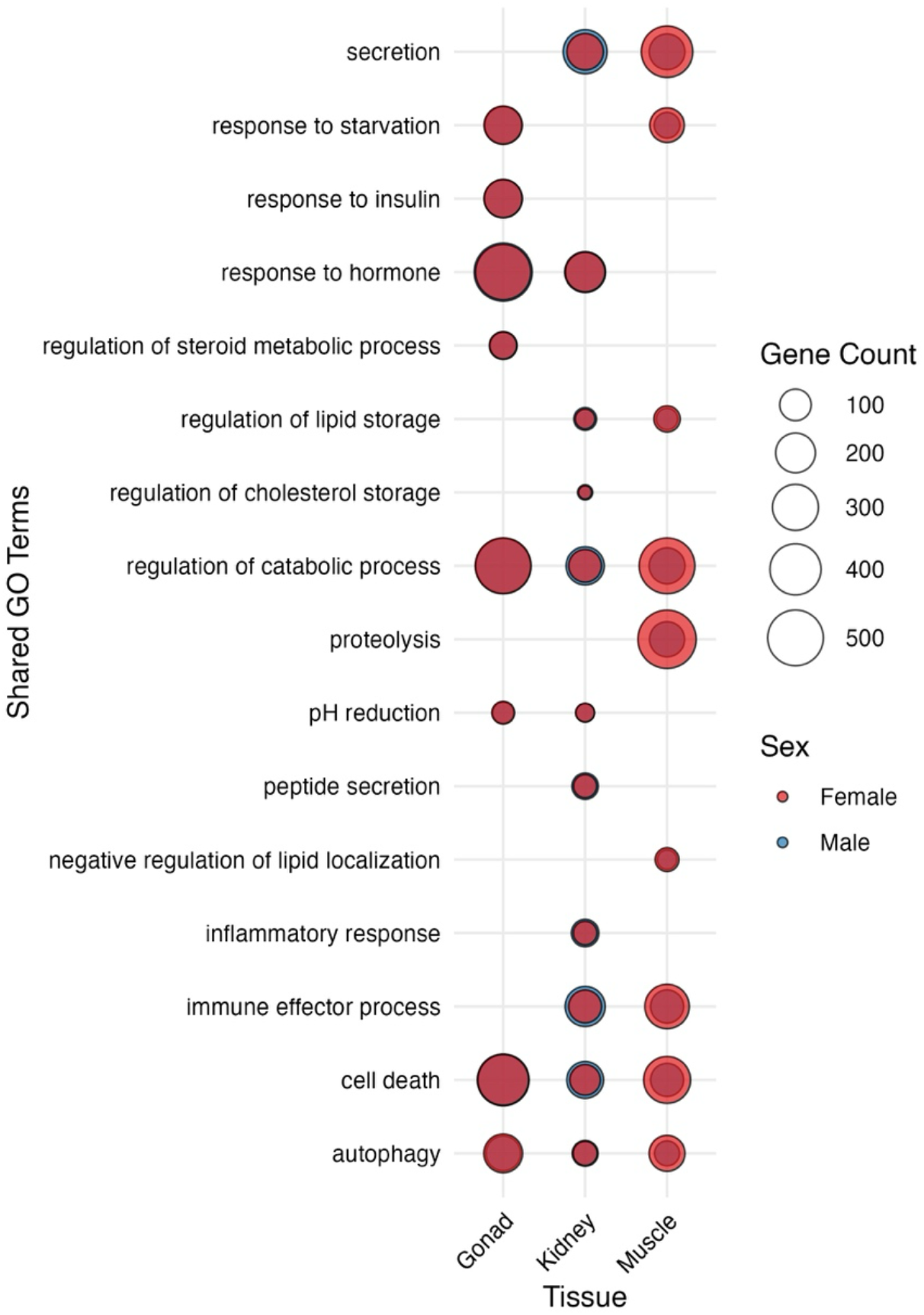
Gene Ontology enrichment of upregulated genes during the transition to the spawning ground. Superimposed bubble plots depicting gene involvement in significantly enriched biological pathways between sexes. The x-axis indicates tissue type, while the y-axis lists shared GO terms. Bubble size is proportional to the number of genes associated with each GO term, and color denotes sex (female, red; male, blue). Overlapping male and female signals within a tissue indicate GO terms shared between sexes, whereas single-color bubbles reflect sex-specific enrichment.

## 3. Results

### 3.1 Patterns of gene regulation during pink salmon spawning maturation

Expression changes from skeletal muscle, head kidney, and gonads of migrating male pink salmon indicated significant molecular changes during the final stage of the migration, progressing through spawning (Figure 2A, Supplemental Table 1). Skeletal muscle had an increase from 301 to 1,499 upregulated genes, and 361 to 3,722 downregulated genes in the transition from ocean to river and river to spawning, respectively. During the same period, the head kidney increased from 91 to 2,272 upregulated genes, and 52 to 3,078 downregulated genes (Figure 2A). The testes had the greatest number of differentially expressed genes during the ocean to river to spawning transition, changing from 5,862 to 10,222 upregulated genes, and 5,017 to 13,431 downregulated genes (Figure 2A). In females, skeletal muscle saw increases in the number of differentially expressed transcripts between the ocean to river to spawning transition (501 to 4,927 upregulated genes, and 507 to 2,677 downregulated genes; Figure 2A). Similarly, the head kidney changed from 1,059 to 1,185 upregulated genes and from 929 to 1,339 downregulated genes. The ovaries of spawning pink salmon increased gene expression from 202 to 5,564 upregulated genes, and 218 to 5,846 downregulated genes.

In males, there were 1,449, 2,236, and 6,871 genes that were only upregulated in the final stages of maturation (no change between ocean and river and changed during spawning), in skeletal muscle, head kidney, and testes, respectively (Figure 2B, Supplemental Table 1). Females had 4,725 skeletal muscle, 1,060 head kidney, and 5,485 ovary genes upregulated during spawning (Figure 2B). In male skeletal muscle, head kidney, and gonads there were 3,636, 3,044, and 8,138 genes downregulated, and females had 4,302, 1,009, 5,676 downregulated genes, during the final transition from the river to the spawning ground (Figure 2B, Supplemental Table 1).

### 3.2 Gene ontology highlights biological pathways integral to spawning phenotypes in pink salmon

We conducted a comprehensive gene ontology (GO) enrichment analysis on filtered genes that were upregulated during the transition from the river to the spawning ground to identify shared biological processes across sexes and tissues (Figure 3). Skeletal muscle processes that were shared across sexes included an upregulation of genes involved in metabolic processes, such as increased starvation response (padj. = 0.0045 male, 0.001 female), changes in lipid storage (pad.= 0.0032 male, 0.0462 female), and the negative regulation of lipid localization (padj. = 0.0126 male, 0.0412 female). Tissue degenerative processes were also increased in both sexes, such as the regulation of tissue catabolism (padj. = 1.88E-05 male, 6.78E-30 female), proteolysis (padj. = 0.0001 male, 6.84E-49 female), autophagy (padj. = 0.0003 male, 3.41E-19 female), and cell death (padj. = 0.0017 male, 0.00583817 female). There was also an upregulation of immune effector processes (pad. = 1.88E-05 male, 0.037917544 female) in skeletal muscle during spawning (Figure 3).

Biological processes upregulated in the head kidney of both males and females included changes in the regulation of lipid (padj. = 0.0134 male, 0.0149 female) and cholesterol storage (padj. = 0.0048 male, 0.0004 female). Unlike skeletal muscle and gonads, the head kidney did exhibit a starvation response in either sex. Like skeletal muscle, there was an increase in tissue catabolism (padj. = 0.0114 male, 2.15E-05 female), autophagy (padj. = 0.0113 male, 1.93E-06 female), and cell death (padj. = 0.0069 male, 0.002 female). Immune effector processes (padj. = 3.66E-16 male, 7.07E-12 female), and inflammatory responses (padj. = 0.023 male, 0.0117 female) were also upregulated in the head kidney of both sexes, as well as the tissue response to hormones (padj. = 0.0012 male, 0.0018 female; Figure 3).

In the gonads of both sexes there was evidence of an upregulation of genes involved in the response to starvation (padj. = 0.0099548 male, 6.97E-11 female), but no changes in the regulation of lipids were identified. Catabolic processes (padj. = 0.0015 male, 2.99E-19 female), autophagy (padj. = 0.00005 male, 1.44E-31 female), and cell death (padj. = 0.0037 male, 3.69E-08 female) were highly upregulated in the gonads like other tissues. Interestingly, the gonads were the only tissue to exhibit a significant upregulation of the insulin signaling pathway (padj. = 0.0006 male, 01.68E-08 female) . Like the head kidney, the gonads had an upregulation of hormone response pathways (padj. = 6.97E-11 male, 4.12E-06 female). The gonads also had an upregulation of genes involved in regulating steroid metabolism (padj. = 0.03065401 male, 0.01278505 female; Figure 3).

### 3.3 Endocrine receptor regulation in spawning salmon depends on ohnolog-specific expression patterns

The thyroid stimulating hormone receptor (*tshr*) was upregulated in the muscle and head kidney of spawning salmon (Figure 4). While minimally expressed in the ovaries, *tshr* was up regulated in the testes during river entry and decreased in expression during spawning. Follicle-stimulating hormone receptor (*fshr*) expression was highly expressed in the gonads with a spike at spawning (Figure 4). Similarly, the luteinizing hormone/choriogonadotropin receptor (*lhcgr*) presented high expression in the gonads with a spike at spawning, while also being expressed at a much lower level in the head kidney prior to spawning. Insulin receptor beta (*insrb*) had low expression consistently across migration in the muscle, head kidney, and ovaries, but was progressively upregulated during migration in the testes (Figure 4). Glucose transporter type 4 (*glut-4*) was expressed only in the muscle and spiked at spawning, while glucose transporter type 1 (*glut-1*) was expressed in all tissues, increasing in expression at spawning. We observed ohnolog-specific expression changes among growth hormone receptor genes. The growth hormone receptor alpha (*ghra)* had low or no expression in any of the tissues analyzed, growth hormone receptor like (*ghr-like*) had modest expression in most tissues and migratory stages except for in the ovaries prior to spawning (Figure 4). Growth hormone receptor beta subunit (*ghrb)* was highly expressed across all tissues and spiked at spawning (Figure 4). Glucocorticoid receptor ohnologs showed divergent expression across tissues and sexes. Receptors *gr* and *gr-like 696* (LOC123993696) were consistently elevated, while glucocorticoid receptors *gr-like 756* (LOC123998756) and *gr-like 423* (LOC124011423) showed low expression across all tissues. Estrogen receptor 1 (*esr1*) had elevated expression in the ovaries, low expression in the head kidney and female muscle, and a spike in male muscle and gonads at spawning (Figure 4). Estrogen receptor 2 subunit alpha (*esr2a*) showed consistently low expression in the muscle and head kidney, but high expression in the gonads, particularly in females. Estrogen receptor subunit beta (*esrrb*) presented mild expression in the head kidney, with low or no expression in the gonads and muscle (Figure 4). Two androgen receptor ohnologs, *ar-like 663* (LOC123993663) and *ar-like 430* (LOC124011430) presented elevated expression across all tissues, with particularly high expression of *ar-like 663* found in the muscle and male gonads. Androgen receptor *ar-like 847* (LOC124008847) was lowly, or unexpressed, across all tissues except male gonads, while the ohnolog *ar-like 303* (LOC123999303) was unobserved in the head kidney, consistently low in the gonads, and minimally up-regulated in the muscle across the migration.

**Figure 4:**
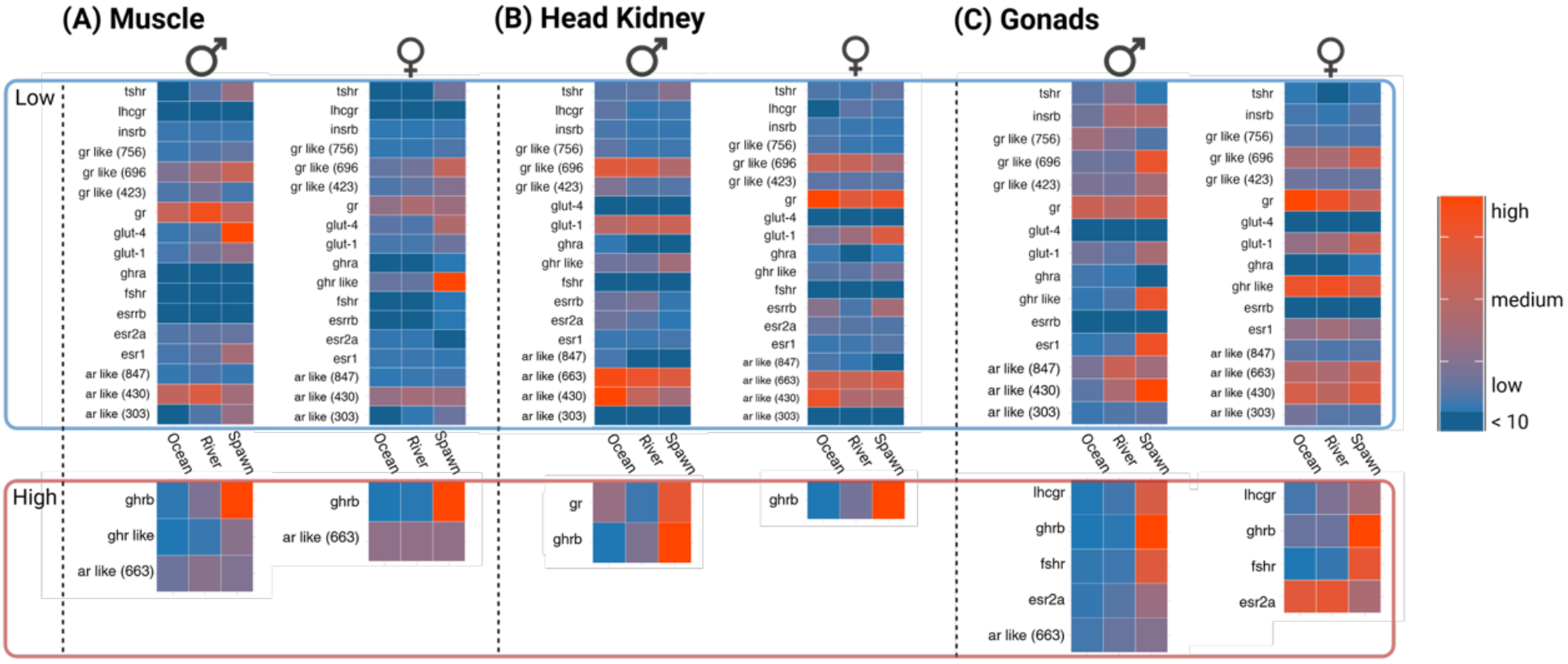
Expression of key endocrine receptors by sex and tissue across migration. Heatmaps depict normalized expression levels of endocrine-related receptors and signaling genes across ocean, river and spawn, shown separately for males (♂) and females (♀) by tissue. Expression values are scaled within tissues and visualized using a blue–red color gradient, where blue indicates low relative expression and red indicates high relative expression. The upper panel (outlined in blue) highlights low expression genes, whereas the lower panel (outlined in red) shows a subset of receptors with high expression. Gene ohnologues are demarked with the last 3 digits of their ncbi gene number annotation.

## 4. Discussion

While the physiological changes that occur during the spawning migration of Pacific salmon have been examined, there is relatively little known about the molecular mechanisms underlying these processes. The results of our study provide molecular evidence for physiological processes ongoing in semelparous pink salmon (22). Our transcriptional analysis revealed a spike in differentially regulated genes within the gonads, head kidney, and skeletal muscle of pink salmon transitioning from the river to the spawning ground, likely coding for late-stage development of maturing gametes and secondary sex traits. The length of salmon migrations and the time they spend in freshwater vary widely between species and populations. Salmon that have longer migrations to the spawning site or enter the river early (e.g., spring run Chinook) retain optimal body condition longer prior to spawning. This suggests that internalized migration timing, not the external stimuli of saltwater to freshwater transition, drives the physiological changes observed during a salmon’s migration. Our transcriptomic analysis reflects this observation and provides evidence of which genes are involved in this final maturation process. The spike in differentially regulated genes appears to coincide with the peak serum levels of stress and sex steroid hormones, further implicating them as a major regulator for spawning phenotypes (29, 30).

As expected, we observed the most differentially regulated genes in the gonads, which is common during reproductive maturation in other species, as the gonads are a transcriptionally active tissue (31). Skeletal muscle and head kidney share a similar rise in gene expression during the transition from the river to the spawning ground. A greater number of differentially regulated genes was observed in the female skeletal muscle, but this observation was reversed in the head kidney, with males having more differentially regulated genes. This could reflect distinct physiological differences between the sexes in these tissues or simply the stage of the maturation process when the sexes were sampled. Likely, it is a combination of both. It is expected that the sexes likely differ in the number of genes necessary to control sex-specific processes or the response to sex hormones. It is also true that males tend to arrive earlier than females and may be farther along the maturation process at the time of sampling, which may be reflected in the transcriptomic data.

Gene ontology analysis found that regulation of catabolism and tissue degeneration processes (autophagy and cell death processes) were significantly up-regulated in spawning pink salmon. While this was expected for skeletal muscle and the head kidney, we were surprised that the gonads exhibited expression signatures of catabolic regulatory processes at the time of spawning. Immune effector processes, which can be associated with inflammation and tissue degeneration, were upregulated in the kidneys and skeletal muscle but not in the gonads. This could hint at the type of tissue catabolism occurring in the tissues, with the catabolic signal derived from the gonads possibly reflecting the physiological changes occurring during final maturation of the gonads (32, 33, 34, 35).

Migrating salmon do not feed, and the impacts of this were observed in the gonads and skeletal muscle. Interestingly, the head kidney did not show the transcriptional signature associated with starvation in either sex. Physiological evidence suggests that tissue degeneration in spawning salmon is driven by a combination of two prominent stimuli, hypercorticotropism in spawning fish that promotes an overactive stress response, and/or the depletion of tissue-specific energy reserves (18, 36, 37). The lack of a starvation response in the head kidney contrasts expectations of the deteriorative processes in salmon and suggests that the head kidney may be able to obtain energy for adrenal hyperplasia and steroidogenesis (6, 18) despite the lack of nutrition in other tissues. This is supported by the observed expression changes in genes regulating lipid homeostasis observed in the head kidney and skeletal muscle, and the negative localization of lipid in the skeletal muscle (Figure 3). This mobilization of lipids was not observed in the gonads of either sex. This may suggest that lipids are being mobilized in tissues like the head kidney and skeletal muscle, and blood sugar is being used to fuel gonad function during the late stage of spawning (19, 38). Interestingly, while we assumed skeletal muscle would be a sink for hormone signaling, this was not the case, and the head kidney and gonads were the only of the three tissues to exhibit transcriptional changes that were involved in hormone responses. This could suggest that skeletal muscle is in full catabolic decline in spawning salmon and is acting primarily as an energy source during this final life stage.

To gain a deeper understanding of the endocrine pathways influencing the gonads, head kidney, and skeletal muscle of pink salmon, we cross-referenced our transcriptome data with previously published hormone profiles occurring during salmon maturation (20, 31, 32, 41-44). This analysis included a comparison of the expression pattern of different endocrine receptor ohnologs. Salmonids have experienced two evolutionary whole genome duplication events, and approximately half of the duplicated genes have been retained in the salmonid genome (39, 40). Whether these ohnologs have conserved or divergent expression patterns or function across salmon tissues, is currently unknown.

Thyroid stimulating hormone receptor levels were up-regulated in skeletal muscle and head kidney during spawning, but not in the gonads, despite the fact that thyroid hormone levels decline at spawning (Figure 4) (29, 41, 42). The increased expression of *tshr* may reflect a compensatory mechanism that is commonly observed in tissues, a feedback response that is believed to increase sensitivity to lower levels of thyroid hormones (29, 41, 42). Growth hormone receptor alpha (*ghra*) exhibited no distinct expression changes across migration (low-expressing tissues), while growth hormone receptor beta (*ghrb*) was highly expressed and spiked at spawning across all tissues, indicating that *ghrb* is likely the primary mediator of growth hormone responses during spawning (Figure 4) (43–46). Despite apparent intolerance to glucose, glucose transporter 1 was expressed at a relatively constant level throughout the migration while showing modest increases at spawning across all tissues (Figure 4) (47–49). Alternatively, glucose transporter 4 expression was in skeletal muscle spiking at the time of spawning.

The gonadotropin receptors, follicle-stimulating hormone receptor (*fshr*) and luteinizing hormone receptor (*lhcgr*), help orchestrate reproductive maturation and we observed low baseline levels of these receptors prior to spawning and then a significant up-regulation in the gonads of both sexes at spawning (30, 50, 51). This highlights the fact that low gonadotropin receptor expression is necessary for gonadal development in the ocean and river, while elevation at spawning reflects the pulse of gonadotropins that accompanies the final maturation of the gonads (30, 31, 51–53).

In female salmon, estrogen is generally elevated until spawning, when it drops prior to death (30, 54), but not found in males and confirmed by total loss of aromatase expression (5, 55). Previous studies have shown that introduction of estrogen into male salmon elicits notable morphological changes similar to females (5). Receptors *esr1* and *esr2a* were expressed in the gonads of both sexes, with *esr2a* being the likely dominant mediator of estrogen signaling in the gonads based on its higher level of expression (Figure 4). The expression of *esr2a* in the ovaries decreased at spawning, following published trends in estrogen levels in spawning salmon (30, 50), while *esr2a* was mildly upregulated at spawning in males. The expression of estrogen-related receptor beta (*esrrb*) showed elevated expression in the head kidney during the early stages of the migration, but its expression level was relatively modest, and it role in this tissue is unclear. We identified expression of all 4 duplicated copies of androgen receptor ohnologs in pink salmon, which had differential expression across tissues. Testosterone trends are known to rise progressively throughout migration and spike at spawning in males and females, changing gene expression to manifest conserved and divergent secondary sex characteristics (5, 53). Only the expression of *ar like 430*, the most broadly expressed androgen receptor ohnolog, matched this increasing pattern of expression in the testes (Figure 4). In the tissues analyzed, it was clear that there is specialized roles for androgen receptor ohnologs across tissues with dominant forms of androgen receptors being expressed in muscle (*ar like 430*), head kidney (*ar like 663* and *ar like 430*), and the gonads (Testes: *ar like 847, ar like 430*; Ovaries: *ar like 663, ar like 430*). The differential expression of androgen receptors may help explain sexually dimorphic phenotypes between sexes and highlight the importance of testosterone in the sexual maturation of both male and female salmon.

Adrenal hyperplasia within the head kidney is believed to be stimulated by sex steroids, resulting in elevated cortisol levels that rise steadily through to spawning (6, 20, 56). Similar to estrogen receptors and androgen receptors, glucocorticoid receptors are ligand dependent nuclear receptors that can act as chaperones to other transcription factors (57). Evidence shows that given a proper environment, glucocorticoid complexes will even heterodimerize with other ligated complexes, including androgen receptors (58). Of the four glucocorticoid receptor ohnologs in pink salmon, *gr* dominated expression across tissues and sexes. The *gr like 696* ohnolog was also expressed at strong levels in all tissues with a decrease in expression at spawning in the head kidney and an increase in skeletal muscle and gonads (Figure 4). This is evidence for ubiquitous influence of the stress hormone coopting the function of sex steroids to help ensure spawning success at the risk of senescence.

Together, our multi-tissue transcriptomic analysis shows that the most dramatic molecular remodeling in pink salmon occurs at spawning rather than during the saltwater-to-freshwater transition, supporting the idea that spawn timing promotes the maturation program in Pacific salmon. Across gonads, head kidney, and skeletal muscle, we identified coordinated activation of catabolic and degenerative pathways (autophagy, proteolysis, cell death) with strong immune signaling in somatic tissues and evidence of lipid mobilization. Finally, endocrine receptor profiles revealed ohnolog-dominant signaling patterns that likely shape key hormonal responses during semelparous maturation and senescence.

### Perspectives and Significance

This study establishes the landscape of transcriptomic activity across the saltwater to freshwater migration of semelparous salmon. The findings implicate several biological processes and endocrine mechanisms that may bolster the physiological changes observed in Pacific salmon prior to their death.

## DATA AVAILABILITY

Source data for this study are openly available at NCBI GEO.

All data and code needed to replicate the results of this study are available on github.

## SUPPLEMENTAL MATERIAL

Supplemental Table 1

## ACKNOWLEDGMENTS

The authors would like to thank the help of Dr. Brad Dimos for guidance on the analysis of the transcriptome data. For their contribution during sample collection, we would like to thank Alex Lopez, Alexander Iritani, Marcus Webb, Tholen Blasko, and Michael Butensky. Special thanks to the Blasko family for private access to the Skykomish River during sampling.

## GRANTS

The research was supported by funding from the United States Department of Agriculture, National Institute of Food and Agriculture grant [2021-67015-33400] and the Foundation for Food and Agricultural Research (FFAR) on competitive grant #FF-NIA21—0000000050.

## DISCLOSURES

The authors have no disclosures to make.

## AUTHOR CONTRIBUTIONS

The research questions and experiments were designed by M.B. and M.P. The laboratory experiments were performed by M.B. and he also performed data analysis. Both M.B and M.P. contributed to the writing of the manuscript.

## Notes

### Competing Interest Statement

The authors have declared no competing interest.

