## Supplemental Table 1 for "Characterization of transcriptomic profiles underlying gross morphological changes observed in semelparous pink salmon (*Oncorhynchus gorbuscha*)"

| **Male** | Muscle | HeadKidney | Teste |
|---|---|---|---|
| UPUP | 25 | 5 | 2311 |
| UPFL | 246 | 46 | 795 |
| UPDN | 30 | 40 | 2756 |
| FLUP | 1449 | 2236 | 6871 |
| FLFL | 15131 | 21721 | 5217 |
| FLDN | 3636 | 3044 | 8138 |
| DNUP | 25 | 31 | 1040 |
| DNFL | 230 | 20 | 1440 |
| DNDN | 106 | 1 | 2537 |
| T1 (up/dn) | 301/361 | 91/52 | 5862/5017 |
| T2 (up/dn) | 1499/3722 | 2272/3078 | 10222/13431 |

| **Female** | Muscle | HeadKidney | Ovary |
|---|---|---|---|
| UPUP | 73 | 27 | 28 |
| UPFL | 225 | 708 | 79 |
| UPDN | 203 | 324 | 95 |
| FLUP | 4725 | 1060 | 5485 |
| FLFL | 11632 | 24324 | 18213 |
| FLDN | 4302 | 1009 | 5676 |
| DNUP | 129 | 98 | 51 |
| DNFL | 206 | 825 | 93 |
| DNDN | 172 | 6 | 74 |
| T1 (up/dn) | 501/507 | 1059/929 | 202/218 |
| T2 (up/dn) | 4927/2677 | 1185/1339 | 5564/5846 |

**Supplemental Table 1. Tables depicting number of genes that filtered into each expression pattern throughout migration by sex.** Filtration of total genes by relative expression (+/- 0 Log2foldchange, p-value $< 0.05$) concatenated across all stages generated 9 groups, 2 of which were extracted for further analysis (FLUP, FLDN).
